# The ulcerative colitis associated gene FUT8 regulates the quantity and quality of secreted mucins

**DOI:** 10.1101/2021.11.30.469270

**Authors:** Gerard Cantero-Recasens, Carla Burballa, Yuki Ohkawa, Tomohiko Fukuda, Yoichiro Harada, IBD Character Consortium, Amy Curwin, Nathalie Brouwers, Gian A. Thun, Jianguo Gu, Ivo Gut, Naoyuki Taniguchi, Vivek Malhotra

**Author notes:** Corresponding authors: Gerard Cantero-Recasens, Vivek Malhotra.

## Abstract

Fucosylation of mucins, the main macrocomponents of the mucus layer that protects the digestive tract from pathogens, increases their viscoelasticity and shear stress resistance. These properties are altered in patients with ulcerative colitis (UC), which is marked by a chronic inflammation of the distal part of the colon. Here we show that the levels of Fucosyltransferase 8 (FUT8) and specific mucins are increased in the distal inflamed colon of UC patients compared to normal individuals. Overexpressing FUT8, as observed in UC, in mucin-producing HT29-18N2 colonic cell line increases trafficking of MUC1 to plasma membrane and secretion of MUC2/MUC5AC. FUT8 depletion (FUT8 KD), instead, causes intracellular accumulation of MUC1 and alters the ratio of secreted MUC2 to MUC5AC. Mucins secreted by FUT8 overexpressing cells are more resistant to shear stress compared to mucins secreted by FUT8 KD cells. These data fit well with the *Fut8*^−/−^ mice phenotype, which are protected against UC. *Fut8*^−/−^ mice exhibit a thinner proximal colon mucus layer with an altered ratio of neutral to acidic mucins compared to *Fut8*^+/+^ mice. Together, these data reveal that FUT8 optimizes the viscoelastic properties of the extracellular mucous by controlling the quantities of mucins secreted.

**SIGNIFICANCE STATEMENT:** Mucins, the major components of the mucous barrier that protects our body from pathogens, are heavily glycosylated proteins. Changes their amounts and properties will alter the viscosity of mucous. Here we show that FUT8, a glycosylation enzyme of the Golgi apparatus, can control the viscosity of secreted mucins. Mucin secreting cells of the distal colon express FUT8, but their levels are altered in Ulcerative colitis patients. As a result, mucous produced by these cells is easily washed away, which exposes them to pathogens. We suggest that a defective mucous production is the main cause of initial inflammation observed in disease. Our findings help in understanding how cells control the quality of mucins and provide a means to prevent Ulcerative colitis.

## INTRODUCTION

The gastrointestinal tract is an optimal habitat for bacteria (high nutrient content, ideal salt concentration and temperature), yet their contact with epithelial cells is limited because of the secreted mucins that compose the mucus layer (1, 2). This layer has two main functions: 1) to protect the epithelium, and 2) to remove pathogens from the body. To date, five gel-forming mucins have been described in humans, including MUC2 as the main mucin for the intestine and colon. Gel-forming mucins are highly glycosylated, which includes fucosylation, during their transit in the Golgi complex (2, 3). These heavily glycosylated mucins are then packed into specialized micrometre sized granules where they can reach molecular weights of up to 50 MDa (4–6). Mature granules fuse to apical plasma membrane by a Ca^2+^ dependent process to release their contents in the absence of extracellular agonist (Baseline Mucin Secretion or BMS) (7) and upon external agonist stimulation (Stimulated Mucin Secretion or SMS) (8). In the extracellular spaces, mucins swell several hundred times their dehydrated volume and together with water, ions and other solutes form the mucus layer. Regulation of mucin secretion ensures the right amounts of mucins depending on the needs of the tissue; while differential mucin glycosylation can modify the rheological properties of the mucus layer for example fucosylation affects viscoelasticity of the mucus. Therefore, alterations in mucin secretion and properties can affect the quality of mucus layer, thus facilitating bacterial invasion and inflammation (9–11).

Inflammatory Bowel Disease (IBD), which includes Crohn’s disease (CD) and Ulcerative Colitis (UC), is characterized by relapsing inflammation of the gastrointestinal tract. Interestingly, quantity and quality of mucins are affected in both CD and UC, especially affecting the colonic mucus layer (12, 13). In healthy colon, the mucus forms two layers: A) an outer layer (less dense and non-attached) colonized by bacteria; and B) an inner layer (dense and attached to the epithelium) impenetrable to bacteria (1, 11). However, in ulcerative colitis patients the colonic mucus layer is thinner and more penetrable to fluorescent beads, and glycosylation correlate with the severity of inflammation (14–16). Importantly, the MUC2 knockout (KO) mice develop spontaneous colitis, revealing the critical role of mucins in UC (17). Several genetic factors have been identified during the last years for UC (18), amongst them Fucosyltransferase 8 (FUT8) (19, 20), which is the only enzyme responsible for the α1-6-linked fucosylation that adds fucose to the innermost GlcNAc residue of an N-linked glycan (21). It is important to note that mucin fucosylation increases mucus viscoelasticity (higher resistance to shear stress and deformation) (22, 23) and, interestingly, we identified FUT8 as a putative regulator of MUC5AC secretion in a genome wide screen (24). In addition, previous studies showed that *Fut8*^−/−^ KO mice develop less-severe colitis than *Fut8*^+/+^ wild type (WT) mice (19).

Clearly, mucus properties and FUT8 levels are altered in UC, which raises the question whether FUT8 controls trafficking of mucins and the viscoelastic properties of secreted mucins? Why is the lack of FUT8 protective against UC? In this study we have used colonic cell lines, in-house gene expression data of colon biopsies from IBD cases, and *Fut8*^−/−^ mice to investigate the significance and role of FUT8 in UC. We have found that FUT8 is required for the correct sorting and trafficking of transmembrane and secreted mucins, as well as for normal viscoelastic properties of mucins. In addition, deletion of FUT8 alters acidic/neutral mucins ratio in the colonic mucus layer of mice, which likely affects the inflammatory response and protects against the pathology. The description of our data follows.

## RESULTS

### FUT8 expression is upregulated in inflammatory bowel diseases

Fucosyltransferase 8 (FUT8) is widely expressed in the gastrointestinal tract, but it is particularly enriched in colonic goblet cells (**Supplementary Table 1**, data extracted from Tabula Muris (25)). Next, we ran differential expression analysis using in-house unpublished RNA-seq data from 93 biopsies taken from different intestinal locations (L1, ileum; L2, ascending colon; L3, descending colon; L4, distal colon) of Ulcerative colitis (UC) patients (N=24), Crohn’s disease (CD) patients (N=22) and control subjects without inflammatory bowel disease (IBD) diagnosis (N=16). Our genetic analysis using these RNA-seq data revealed higher FUT8 levels in the proximal colon compared to distal colon of control subjects (3.3-fold higher in L1 compared to L4, p=9.41E-05, 2.2-fold higher in L2 compared to L4, p=0.006), a compartmentation that is lost in inflamed colon of Ulcerative Colitis (UC) patients (L2 vs. L4 Fold change= 1.08, p=0.78), and partially in Crohn’s disease (CD) patients (L1 vs. L4 Fold change=1.8 p=0.004, L2 vs. L4 Fold change=2.1 p=0.006) (**Figure 1A**). Our analysis also showed that RNA levels of several mucins (i.e. MUC1, MUC4, MUC5B, MUC12 and MUC20) are higher in the distal colon compared to proximal colon. This distribution was generally maintained in inflammation (CD and UC). However, some mucins showed inconsistent differential expression between colonic regions, possibly because of low expression levels in general. Our results also show that FUT8 is increased in inflamed colon of CD and UC patients compared to non-inflamed colons, especially in UC (3-fold increase in UC p=9.7E-07, 2.4-fold increase in CD p=5.5E-05). Levels of MUC1, MUC14, MUC16, MUC5AC and MUC5B were suggestively also higher in inflammation whereas MUC12, MUC17 and MUC6 levels are decreased (**Figure 1B**). Statistical significance in these comparisons could not be emphatically confirmed because of the generally small sample size. We therefore suggest that this is considered rather a general trend in changes in the expression of mucins.

**Table 1.**
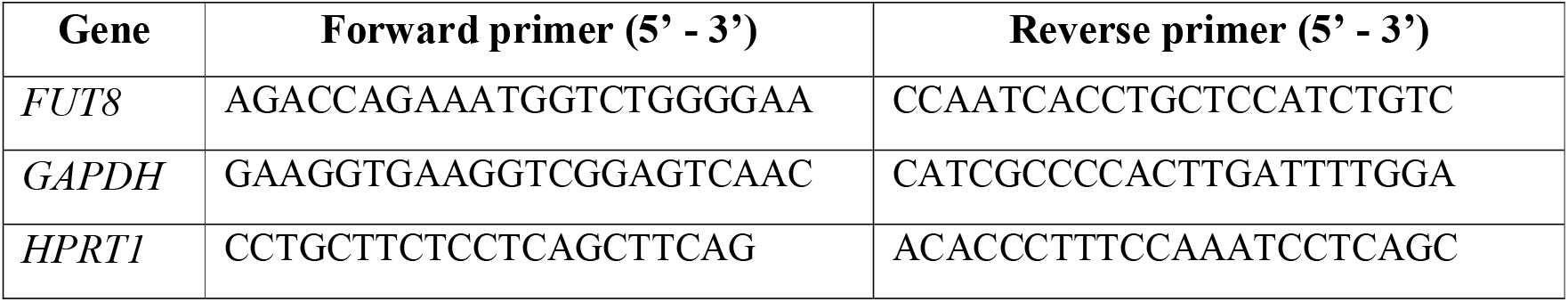
Primer sequences used for detecting mRNA for the respective genes

**Figure 1.**
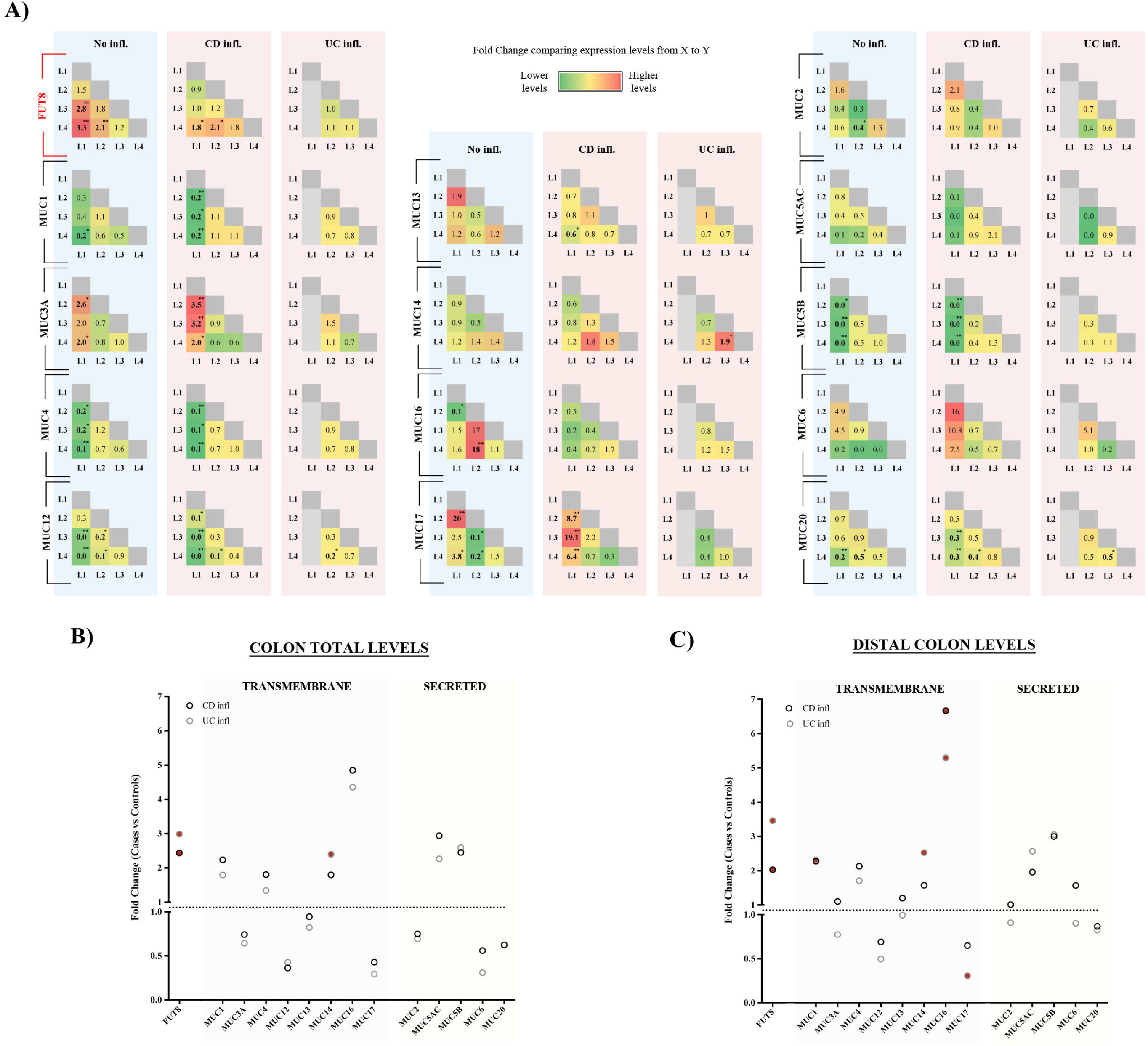
Colonic expression levels of FUT8 and mucins in bowel diseases. (**A)** Expression of FUT8 and mucins was compared between different colonic locations (L1: ileum, L2: ascending colon, L3: descending colon, L4: distal colon) in control subjects, CD patients and UC patients. Data are shown as fold change between locations of the X-axis vs. the Y-axis. **(B)** Total levels of FUT8 and 14 mucins at the colon comparing control subjects *vs.* patients. Data represent the fold change between controls and UC patients (black circles) or CD patients (grey circles). Red-filled circles represent those statistically significant differences (p<0.004). **(C)** Specific levels of FUT8 and 14 mucins at the distal colon comparing the data from patients (UC, black circles; CD, grey circles) with the expression levels of control subjects. Red-filled circles represent those statistically significant differences (p<0.004). Abbreviations: No infl.: Not inflamed, CD infl.: Crohn’s disease inflamed colon, UC infl.: Ulcerative colitis inflamed colon. * p<0.05, **p<0.01

FUT8 has been associated with UC (18) and it is known that the mucus layer of UC patients is thinner and inflammation affects mainly the distal part of the colon (L4 in our analysis). The mucus layer is thicker in CD and can affect any part of the gastrointestinal tract, but more usually the small intestine and proximal colon (26, 27). Accordingly, our expression data showed that FUT8 levels are increased in inflamed distal colon (L4) of UC patients (3.5-fold increase, p=7.14E-08) and also, although at lesser extent, in CD patients (2-fold increase, p=0.004) (**Figure 1C**). Amongst mucins, MUC1, MUC14 and MUC16 levels were increased in UC inflamed distal colon (2-fold, 2.5-fold and 5-fold increase, respectively, p<0.05), while MUC17 levels were reduced (70% reduction, p=0.01). Although not approaching statistical significance, both MUC5AC and MUC5B levels show a tendency to be higher in inflamed UC distal colon (2.5 and 3-fold increase, respectively).

Altogether, our results show that FUT8 levels are increased in inflamed UC distal colon and show a similar trend for several transmembrane (e.g. MUC1) and secreted mucins (e.g. MUC5AC).

### FUT8 regulates export of MUC1, MUC2 and MUC5AC

We identified FUT8 as a putative gene required for mucin secretion triggered in cells treated with Phorbol 12-myristate 13-acetate (PMA) (24). To understand the net effect of FUT8 on mucin secretion and its link to the physiology of mucins in the colon, we used HT29-18N2 cells that can be differentiated into mucin producing cells by serum starvation (28). We generated a stable cell line depleted of FUT8 (FUT8 KD) and a stable cell line that overexpresses FUT8 (FUT8 OV). RNA was extracted from differentiated FUT8 KD and FUT8 OV HT29-18N2 cells and *FUT8* mRNA levels measured by qPCR. Compared to control cells, FUT8 KD cells revealed 70% reduction in *FUT8* mRNA levels, whereas FUT8 OV cells showed 5-fold increase in the cognate mRNA (**Supplementary Figure 1A**), which was also confirmed at protein level by western blot (WB) (**Supplementary Figure 1B**). There was a modest increase of MUC1 levels in cells overexpressing FUT8 (35% increase), but no discernable effect on the levels of secreted mucins MUC2 and MUC5AC (**Supplementary Figure 1C-E**). MUC2 and MUC5AC secretion in absence (Baseline) and presence (Stimulated) of physiological stimulus ATP (100 μM in a solution containing 1.2 mM CaCl_2_) was then examined in these cells. Our results show that depletion of FUT8 increases baseline MUC2 secretion (1.7 fold increase) (**Figure 2A**) and agonist (ATP) induced MUC5AC secretion (1.4 fold increase) compared to control condition (**Figure 2B**). Overexpression of FUT8 increases both MUC2 and MUC5AC baseline and ATP-dependent (stimulated) secretion (MUC2: 1.4 and 1.8 fold increase, MUC5AC: 1.7 and 1.6 fold increase, respectively) (**Figures 2C and 2D**).

**Figure 2.**
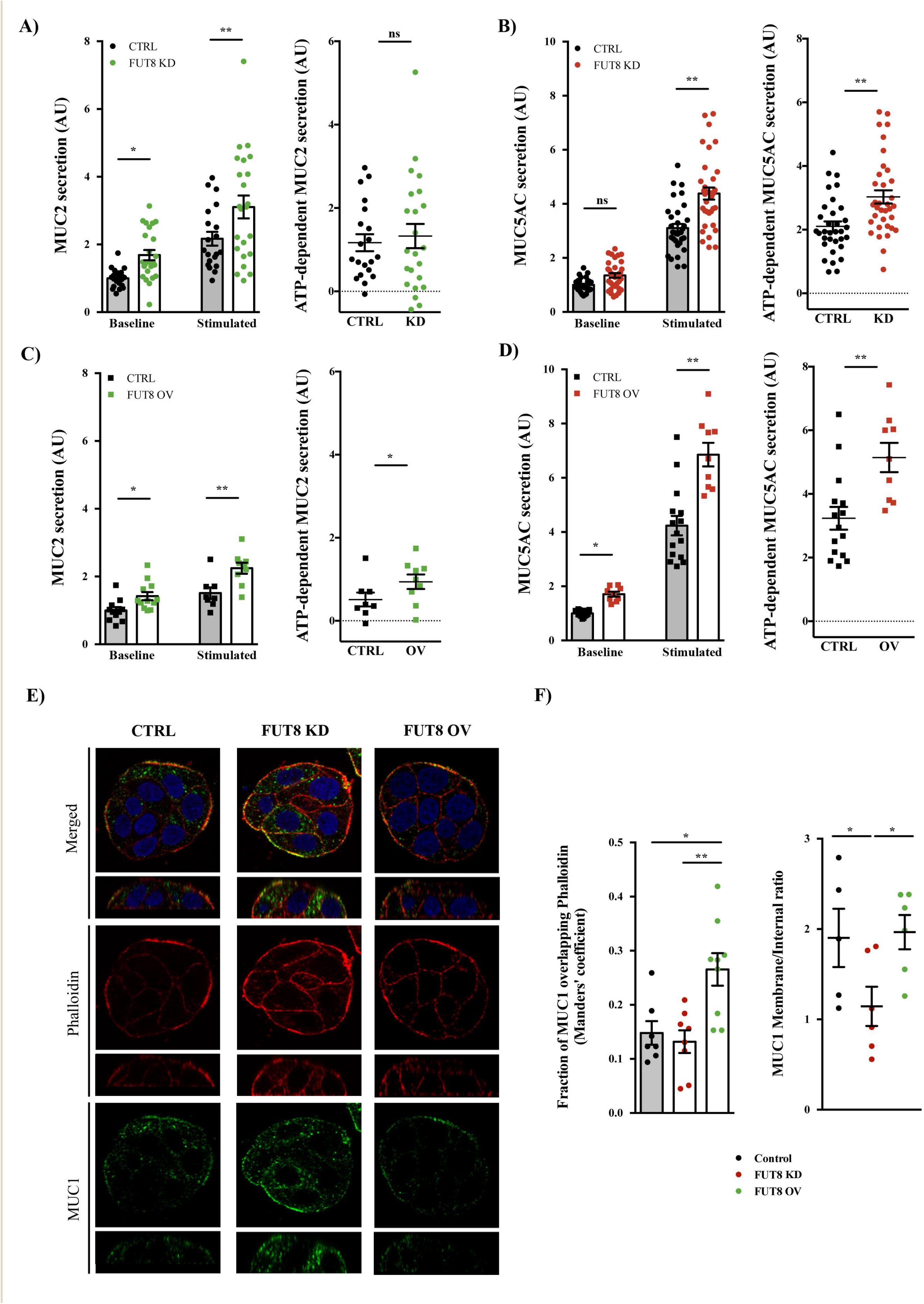
FUT8 levels alter mucin export. **(A and C)** Secreted MUC2 from differentiated control (black circles) and FUT8-KD (green circles) cells or control (black squares) and FUT8-OV (green squares) cells that were incubated for 30 min at 37°C in the absence (Baseline) or presence (Stimulated) of 100 μM ATP. Data were normalized to intracellular actin levels. The y-axis represents normalized values relative to the values of untreated control cells. ATP-dependent MUC2 secretion was calculated from the data in (A) or (B), respectively, as the difference between normalized baseline secretion and stimulated secretion for each condition. **(B and D)** Secreted MUC5AC from differentiated control (black circles) and FUT8-KD (red circles) cells or control (black squares) and FUT8-OV (red squares) cells that were incubated for 30 min at 37°C in the absence (Baseline) or presence (Stimulated) of 100 μM ATP. Data were normalized to intracellular actin levels. The y-axis represents normalized values relative to the values of untreated control cells. ATP-dependent MUC5AC secretion was calculated from the data in (C) or (D), respectively, as the difference between normalized baseline secretion and stimulated secretion for each condition. **(E)** Immunofluorescence Z-stack single planes of control, FUT8-KD and FUT8-OV differentiated HT29-18N2 cells with anti-MUC1 antibody (green), Phalloidin (red) and DAPI (red). Scale bar = 5 μm. **(F)** Left graph. Colocalization between MUC1 and Phalloidin was calculated from immunofluorescence images by Manders’ coefficient using FIJI. Average values ± SEM are plotted as scatter plot with bar graph. The y-axis represents Manders’ coefficient of the fraction of MUC1 overlapping with Phalloidin. **(F)** Right graph. Quantification of MUC1 levels at the membrane compared to intracellular levels in control (black), FUT8 KD (red) and FUT8-OV (green) cells. Data was calculated from immunofluorescence images using FIJI. Abbreviations: Ctrl: control cells, FUT8 KD: FUT8 depleted cells, FUT8 OV: FUT8 overexpressing cells, KD: FUT8 KD, OV: FUT8 OV. * p<0.05, ** p<0.01.

These data reveal that FUT8 levels have different effects on MUC2 and MUC5AC secretion. When presented as the ratio of MUC2 to MUC5AC levels, FUT8 KD cells present higher baseline MUC2/MUC5AC ratio (Ratio=1.35, more MUC2 secretion) but lower stimulated MUC2/MUC5AC ratio (Ratio=0.6, more MUC5AC secretion) than control cells (**Supplementary Figure 2A**). Overexpression of FUT8 has no obvious effect on baseline ratio (Ratio=0.9), but has the opposite phenotype on stimulated MUC2/MUC5AC ratio (Ratio=1.4, more MUC2 secretion) (**Supplementary Figure 2B**). These changes may affect the mucus layer properties due to different MUC2 and MUC5AC fibres formation (1).

Finally, we also studied the effect of FUT8 on MUC1 (a transmembrane mucin). Although MUC1 was transported to the plasma membrane in all cell lines (**Figure 2E**), this was significantly enhanced in FUT8 overexpressing cells (Manders’ coefficient MUC1 vs. Phalloidin Ctrl=0.14, FUT8 KD=0.13 and FUT8 OV=0.27) (**Figure 2F, left panel**). Interestingly, depletion of FUT8 caused intracellular MUC1 accumulation compared to control cells and in cells overexpressing FUT8 (**Figure 2F, right panel**).

### The composition of mucus layer is altered in *Fut8*^−/−^ mice

FUT8 knockout (KO) mice (*Fut8*^−/−^ mice) are less susceptible to ulcerative colitis (19). Previous studies reported lower levels of T-helper 1 and 2 cytokines in *Fut8*^−/−^ mice, which could explain the protective effect in the progression of T-cell mediated colitis (19). However, the effect of FUT8 on initiation of inflammation, which is related to the mucus layer, has not been studied. Based on our data showing that FUT8 controls mucin secretion and trafficking in colon cancer cell lines, we tested the effects of FUT8 deletion on the mucus layer under basal conditions.

Colons from *Fut8*^+/+^, *Fut8*^+/−^ and *Fut8*^−/−^ mice (N=4 per condition, 12-week old) were stained with haematoxylin and eosin (H/E). Histological analysis showed no obvious differences in epithelial hyperplasia, inflammatory infiltrate and oedema at the proximal, middle and distal colon; or in the counting of the mitotic figures between groups (**Supplementary Table 2, Supplementary Figure 3A and 3B**).

We then stained mice colon slices with LCA-lectin, which binds fucosylated proteins (29). Our results confirm that the colonic mucus layer is fucosylated and, importantly, it is reduced in *Fut8*^−/−^ mice (**Supplementary Figure 3C, left panel**). Next, we stained the colonic slices using PhoSL (30), a FUT8-dependent fucosylation specific lectin. Our data revealed a complete absence of PhoSL staining in *Fut8*^−/−^ mice compared to *Fut8*^+/−^ mice (**Supplementary Figure 3C, central panel**), confirming the FUT8-dependent fucosylation of the colonic mucus layer.

To further evaluate the mucus layer in these mice, slices from proximal, medial and distal mice colon (*Fut8*^+/+^, *Fut8*^+/−^, *Fut8*^−/−^) were stained with Periodic acid-Schiff (PAS) and Periodic acid-Schiff-Alcian Blue (PAS-AB) to detect neutral and acidic mucins as described before (7). The thickness of the mucus layer detected by PAS staining (neutral mucins) was not significantly different between conditions at any part of the colon (**Figure 3A-C, upper panels**). However, there was a reduction in the mucus thickness detected by PAS-AB (acidic and neutral mucins) at proximal colon in *Fut8*^+/−^ and *Fut8*^−/−^ compared to *Fut8*^+/+^ mice (12.93 μm and 11.27 μm vs. 23.81 μm, respectively). The mucus thickness was close to the normal in medial (21.27 μm and 24.6 μm vs. 29.02 μm) and distal (22.2 μm and 22.4 μm vs. 23.5μm) colon (**Figure 3A-C, lower panels**). The ratio of PAS-AB to PAS staining decreased from proximal to distal in *Fut8*^+/+^ mice (proximal = 1.8, medial = 1.4, distal = 0.8, proximal *vs.* distal p=0.019), which was not observed in both FUT8^+/−^ (proximal = 0.9, medial = 1.1, distal = 1.3, proximal *vs.* distal n.s.) and FUT8^−/−^ (proximal = 1.0, medial = 1.0, distal = 1.1, proximal *vs.* distal n.s.) mice (**Supplementary Figure 3D**).

**Figure 3.**
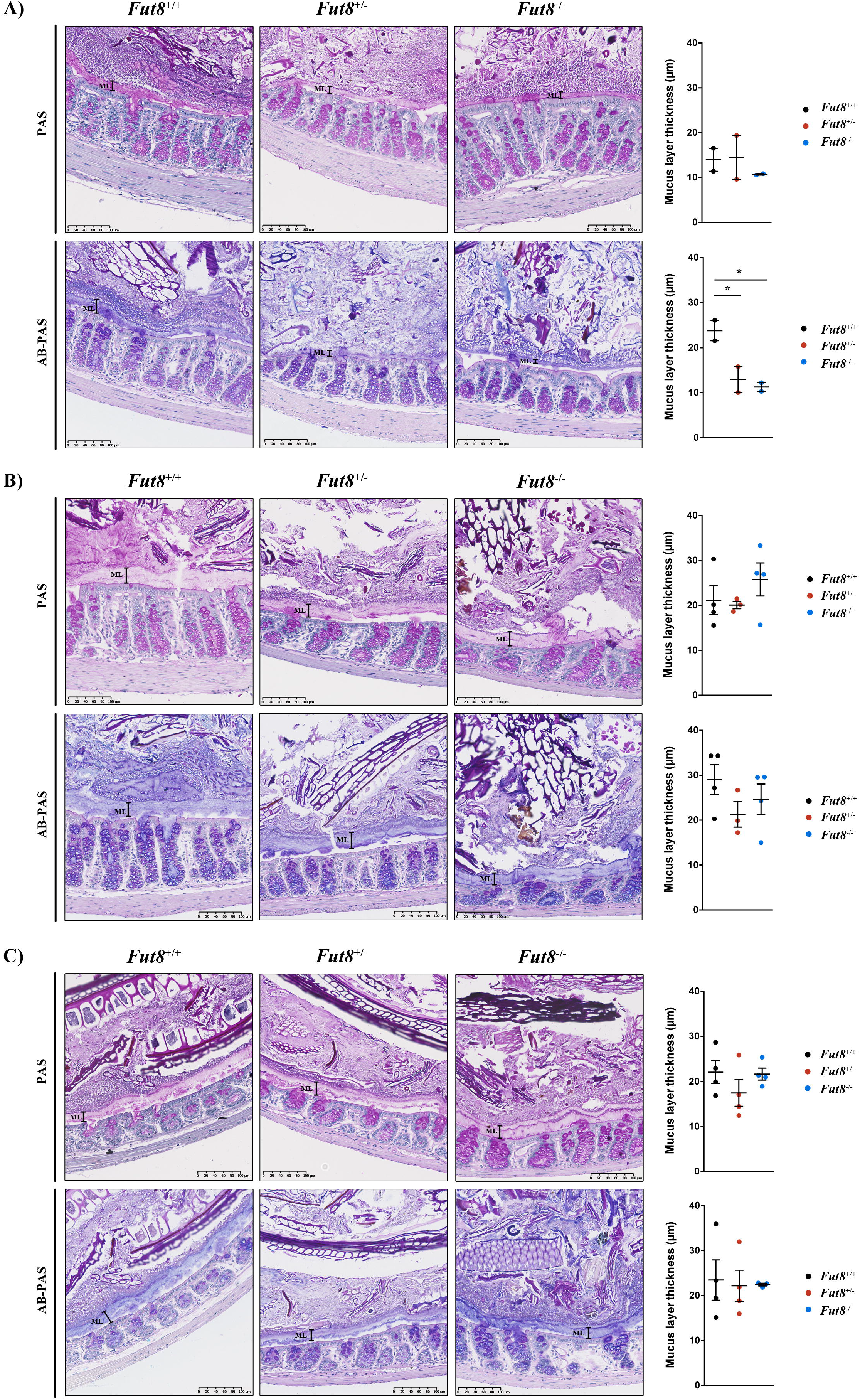
*Fut8*^−/−^ mice present thinner mucus layer at the proximal colon. Representative **(A)** Proximal, **(B)** Medial and **(C)** distal colons of *Fut8*^+/+^ (left panel), *Fut8*^+/−^ (central panel) and *Fut8*^−/−^ (right panel) mice stained with PAS or PAS-AB. Quantification of the mucus layer thickness for each condition is shown next to images. Average values ± SEM are plotted as scatter plot with bar graph (*Fut8*^+/+^: black dots, *Fut8*^+/−^: red dots, *Fut8*^−/−^: blue dots). The y-axis represents the thickness of the mucus layer in μm. *p<0.05.

### FUT8 alters the resistance of mucins to shear stress

Mucin fucosylation affects viscoelastic properties of mucus, increasing its resistance to shear stress. Thus, we tested whether alteration in FUT8 levels affects mucins resistance to shear stress, which would explain why both FUT8 KD and FUT8 OV alter mucin secretion. First, we checked extracellular mucin fibres after stimulation (using 100 μM ATP) in control, FUT8 KD and FUT8 OV cells by immunofluorescence (IF) with and without permeabilization before or after extensive washing with isotonic solution. Unpermeabilized and not washed cells showed extracellular mucin fibres in all conditions. Importantly, MUC5AC fibres formed by FUT8 KD or FUT8 OV cells were different from those secreted by control cells as observed by confocal microscopy (**Figure 4A**). After washing (without permeabilization) few mucin fibres were visible in control or FUT8 KD cells compared to several fibres in FUT8 OV cells (**Figure 4B**). Finally, cells were permeabilised to confirm the presence of mucin granules (**Figure 4C**). This lends support to the hypothesis that in FUT8 OV cells, the secreted mucins are more resistant to shear stress.

**Figure 4.**
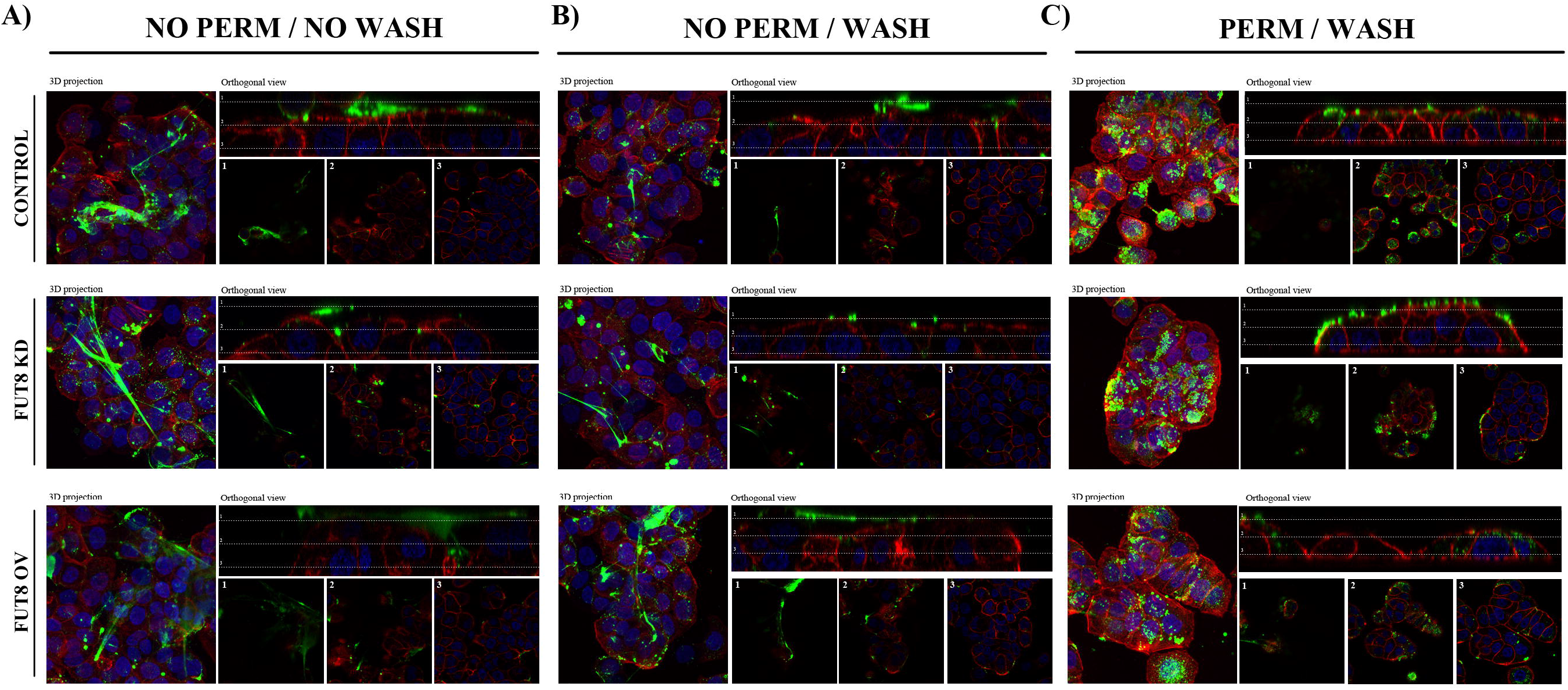
FUT8 affects the shear stress resistance of the mucin fibres. **(A,B)** Control, FUT8 KD and FUT8 OV cells were differentiated for six days and seeded in glass-bottom dishes. After stimulation using 100 μM ATP, cells were processed for immunofluorescence **(A)** before (NO PERM / NO WASH) or **(B)** after extensive washing with isotonic solution (NO PERM / WASH). Cells were stained with anti-MUC5AC (Green), Phalloidin (Red) and DAPI (Blue). In the orthogonal view, dotted lines across the images demarcate the top surface of the cell. Scale bars correspond to 5 μm. **(C)** Control, FUT8 KD and FUT8 OV cells were permeabilized after extensive washing (PERM / WASH) to reveal intracellular MUC5AC granules. Abbreviations: NO PERM / NO WASH: Not permeabilized cells without washing, NO PERM / WASH: Not permeabilized cells with extensive washing, PERM / WASH: Permeabilized cells after extensive washing.

To further test whether FUT8 alters shear stress resistance of mucins, we measured the levels of MUC5AC recovered after three successive washes with isotonic solution (W0, W1 and W2). As shown above, FUT8 depletion leads to higher ATP-dependent MUC5AC secretion, whereas FUT8 overexpression increases both baseline and stimulated MUC5AC secretion. To compare the percentage of MUC5AC recovered with each wash, MUC5AC levels were normalized to the total amount of mucin recovered after three washes. Our results revealed no major differences between control and FUT8 KD cells in the percentage of MUC5AC recovered in the successive washes. However, in cells overexpressing FUT8, a higher percentage of MUC5AC was detected in the first wash after ATP stimulation (75% in FUT8 OV vs. 61% in control cells, p<0.05) than control and FUT8 KD cells. A similar tendency was observed in the first wash for baseline MUC5AC secretion (73% vs. 63%, p=0.13). Consistently, a lower percentage of MUC5AC was detected in the following wash (W2) in cells overexpressing FUT8 compared to control cells (Baseline: 11% vs. 22%, p<0.05; Stimulated: 14% vs. 27%, p<0.05) (**Supplementary Figure 2C and 2D**).

Altogether, these data show that FUT8 depleted cells secrete more mucins that can be easily removed from cell surface, whereas FUT8 overexpressing cells secrete mucins that are more resistant to shear stress and removal.

## DISCUSSION

Strict control of mucin secretion is essential for the correct formation and function of mucus that protects the intestinal epithelium from inflammation and infection (6). Alterations in the secreted quantities or composition due to aberrant biosynthesis, trafficking or secretion of mucins will affect the rheological properties of the mucus barrier affecting bacteria-epithelium contact and leading to inflammatory intestinal pathologies such as Crohn’s Disease (CD) and Ulcerative Colitis (UC) (31). UC is confined to the distal part of the colon and the rectum, and characterized by mucosal inflammation (32). Interestingly, empty goblet cells (mucin-producing cells) is a morphological characteristic used to assess active UC, which could reflect defects in mucin synthesis or enhanced mucin granules exocytosis (14).

We previously identified Fucosyltransferase 8 (FUT8), the only enzyme responsible for the alpha 1-6-linked fucosylation, as a putative component of mucin secretion pathway (24). Interestingly, increased levels of FUT8 are associated with intestinal diseases like ulcerative colitis and *Fut8*^−/−^ mice are protected against severe colitis. Fujii and colleagues have shown that core fucosylation by FUT8 is required for T cell signalling and production of inflammatory cytokines, which could explain the link between FUT8 and UC progression (19). In the present study, we have described FUT8’s involvement in UC pathogenesis by affecting the secretion of specific mucins that control the viscoelastic properties of the mucous layer.

### The expression of FUT8 is altered in the inflamed colon

Our data show that FUT8 levels are higher at the proximal part of the colon, where the mucus layer is thinner to allow contact between the microbiota and epithelium (33). Importantly, FUT8 levels are upregulated in distal inflamed colon of UC patients compared to control subjects. In addition, mice depleted of FUT8 (*Fut8*^−/−^) present a thinner proximal colonic mucus layer (PAS-AB staining) compared to WT mice (*Fut8*^+/+^) and an altered ratio PAS/PAS-AB that is in accordance with FUT8 expression along the colon (higher levels at proximal, lower at distal). In fact, the only differences between *Fut8*^−/−^ and *Fut8*^+/+^ mice are observed at the proximal colon. This fits with our genetic data showing that FUT8 levels are higher at the proximal colon and therefore expected to reveal more differences.

Based on our data, we postulate that FUT8-dependent fucosylation alters the viscoelastic properties of the mucus layer, which likely enhances the contact between microbiota and epithelia. In pathologic conditions FUT8 is upregulated (as observed in distal colon of UC patients), which likely increases interaction between microbiota and epithelium leading to more inflammation. In contrast, in *Fut8*^−/−^ mice lacking fucosylated mucins, bacteria fail to attach and there is less invasion and inflammation, which prevents the development of ulcerative colitis.

### FUT8 regulates MUC1 trafficking

Our results show that FUT8 has a role in trafficking of transmembrane mucins (i.e. MUC1) to the plasma membrane. Intracellular MUC1 levels in FUT8 KD are higher compared to control cells, whereas FUT8 overexpression enhanced MUC1 localization at the plasma membrane. Interestingly, MUC1 has been described to regulate inflammatory responses to infection (34) and serve as adhesion receptor for enteroaggregative *E. coli* (EAEC) (35) and *H. pylori* (36). It is plausible that increased levels of FUT8 (as seen in UC patients) lead to more MUC1 at the plasma membrane, which act as receptor for bacteria allowing epithelial invasion and inflammation. In contrast, low levels of FUT8 provoke intracellular MUC1 retention and may explain the protection against UC observed in *Fut8*^−/−^ mice.

### FUT8 modulates mucin secretion

In the colonic HT29-18N2 cell line, changes in the FUT8 levels modulate mucin secretion and resistance of mucus to shear stress. Our results reveal that FUT8 deletion increases MUC2 baseline secretion and MUC5AC stimulated secretion while FUT8 overexpression enhances both MUC2 and MUC5AC baseline and stimulated secretion (**Figure 3**). Taking into account that fucosylation increases mucus resistance to shear stress (37), we suggest that FUT8’s effect on mucin secretion is due to alteration in mucins’ properties. Accordingly, our data confirm that FUT8 depleted cells secrete more mucins that are easily removed, but FUT8 overexpressing cells secrete mucins that are more resistant to shear stress and removal (**Figure 4**). This also provides a reasonable explanation why *Fut8*^−/−^ mice present thinner mucus layer at the proximal colon (by PAS-AB staining) while FUT8 KD cell lines show enhanced mucin secretion.

### A model for FUT8 on UC pathogenesis

In sum, we propose the following scheme for the role of FUT8 in the pathogenesis of UC **(Figure 5)**.

A. Normally, the colonic mucus barrier is composed of two layers: A) an outer layer where pathogens and microbiota are trapped; and B) an inner layer free of bacteria (**Figure 5, control**). However, due to genetic or environmental factors the mucus layer is altered (e.g. increased expression of MUC5AC or altered glycosylation) allowing bacterial invasion of the colonic epithelium that can lead to inflammation and finally colitis.
B. <u>Cells with low levels of FUT8</u> (which protects against UC) show increased secretion of mucins lacking fucosylation (**Figure 5, Low FUT8**). This lack of fucosylated mucins produces a more stacked inner layer, but an easily removable outer layer, consequently reducing the amount of bacteria and microbiota, and increasing the shear stress that will trigger more mucin secretion (and increased renewal rate). In addition, lack of fucosylation reduces the receptors for bacteria (e.g. MUC1 or fucosylated mucins), which protects the epithelium against bacterial invasion and, therefore, inflammation.
C. <u>High FUT8 levels</u> (similar to the situation at the proximal compared to the distal colon) associated to UC, lead to a mixed inner/outer layer that is more permeable and sticky (difficult to remove) that traps bacteria, which invade and reach the epithelium (**Figure 5, High FUT8**). Finally, cells with high levels of FUT8 present more receptors for bacteria at the plasma membrane, facilitating the invasion of the intestinal epithelium and inflammation that can lead to UC pathogenesis.

**Figure 5.**
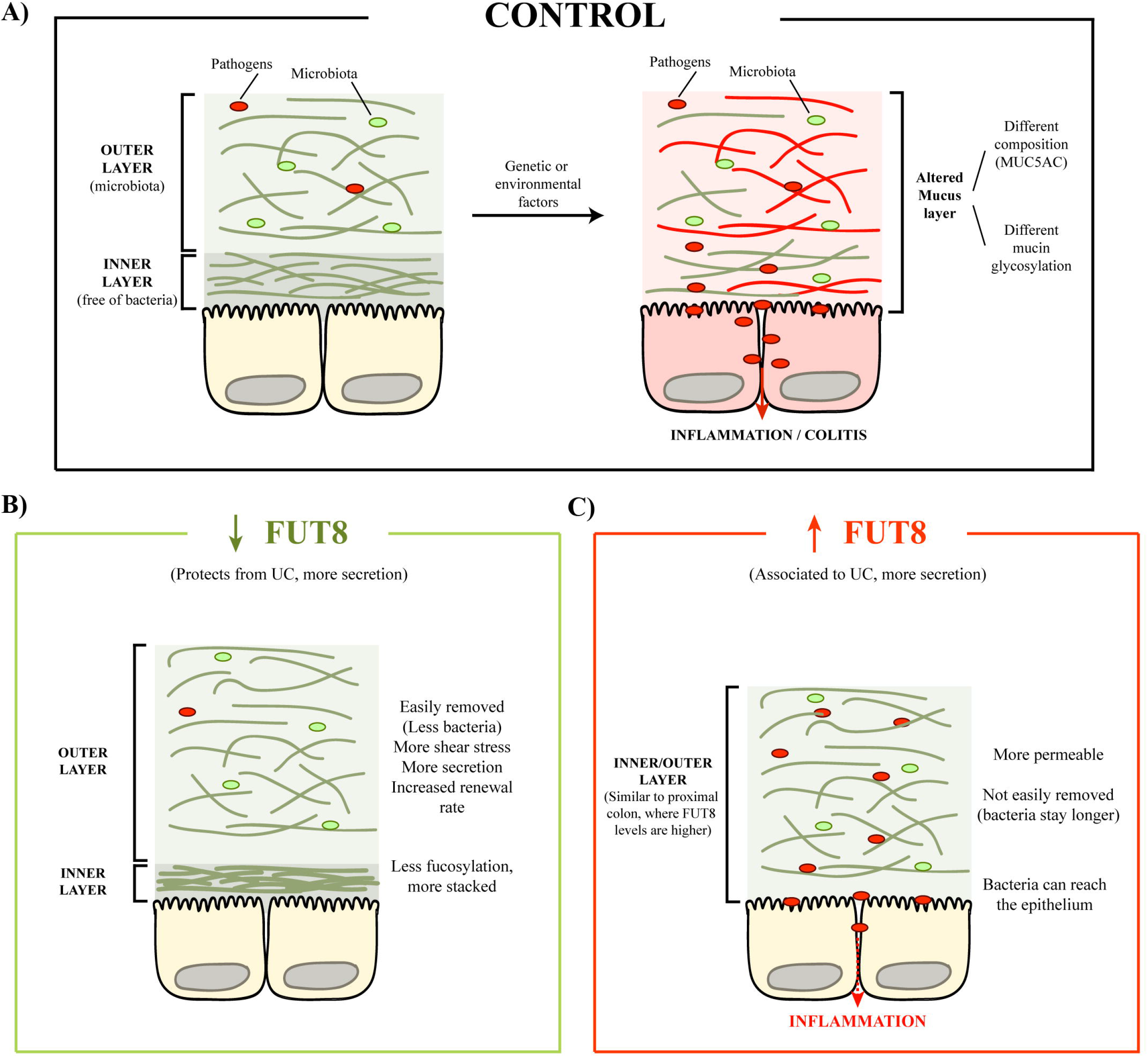
Model for FUT8 role on UC pathogenesis. **(A)** In control cells, both mucus layers are separated in the distal colon, being the pathogens and the microbiota restricted to the outer layer. However, genetic or environmental factors can alter the mucus layer properties, allowing pathogens to reach the colonic epithelium and triggering inflammation. **(B)** In cells with low levels of FUT8, mucins are less resistant to shear stress, which makes the outer layer, together with bacteria, easily removed. In turn, this triggers more secretion, thus increasing the renewal rate and protecting against invasion from pathogens. **(C)** High levels of FUT8, as observed in UC patients, disrupt the mucus layer, making it more permeable to pathogens. In addition, these mucin fibres are more resistant to shear stress, so the bacteria can stay longer and reach the epithelium triggering inflammation. Abbreviations: UC: Ulcerative colitis.

## MATERIALS AND METHODS

### RNA public datasets analysis

The final data set consisted of 62 individuals (24 UC patients, 22 CD patients, 6 healthy controls and 10 symptomatic controls without a IBD diagnosis), from whom up to two biopsies (usually one from an inflamed and one from a non-inflamed location) were available. Locations were categorized into L1 (ileum), L2 (ascending colon), L3 (descending colon), L4 (distal colon). Raw counts were normalized using TMM, which takes library size and RNA-composition into account. Limma-voom was used to run the differential expression analysis. The statistical models were corrected for sex, age and origin (centre). There was no comparison in which two samples from the same individual were included. 19.7k genes showed robust expression across the biopsies. 14 genes encoding different mucins and FUT8 were selected for the present study. Nominal p-values were therefore corrected for multiple testing lowering the significance threshold to 0.05/14=0.004.

### Reagents and antibodies

All chemicals were obtained from Sigma-Aldrich (St. Louis, MO) except anti-MUC2 antibody clone 996/1 (RRID:AB_297837) (Abcam, Cambridge, UK), anti-MUC5AC antibody clone 45M1 (RRID: AB_934745) (Neomarkers, Waltham, MA) and anti-FUT8 antibody (RRID:AB_1523644) (Abcam, Cambridge, UK). Secondary antibodies for immunofluorescence microscopy and dot blotting were from Life Technologies (ThermoFisher Scientific, Waltham, MA, USA).

### Cell lines

HT29-18N2 cells (obtained from ATCC) (RRID:<u>CVCL_5942</u>) were tested for mycoplasma contamination with the Lookout mycoplasma PCR detection kit (Sigma-Aldrich, St. Louis, MO). Mycoplasma negative HT29-18N2 cells were used for the experiments presented here.

### Generation of stable cell lines (shRNA and overexpression)

HEK293T cells (ATCC, negative for mycoplasma) were co-transfected with the plasmid, VSV-G, pPRE (packaging) and REV by Ca^2+^ phosphate to produce lentiviruses. Forty-eight hours post transfection; the secreted lentivirus was collected, filtered and directly added to HT29-18N2 cells. Stably infected HT29-18N2 cells with the different constructs were sorted for GFP signal by FACS.

### Quantitative real-time PCR (RT-qPCR)

Differentiated HT29-18N2 control, FUT8-KD and FUT8 overexpressing (FUT8 OV) cells were lysed and total RNA extracted with the RNeasy extraction kit (Qiagen, Netherlands). cDNA was synthesized with Superscript III (Invitrogen). Primers for each gene (sequence shown below, Table 1) were designed using Primer-BLAST (NCBI) (38), limiting the target size to 150 bp and the annealing temperature to 60°C. To determine expression levels of the different genes, quantitative real-time PCR was performed with Light Cycler 480 SYBR Green I Master (Roche, Switzerland) according to manufacturer’s instructions.

### Differentiation of HT29-18N2 cells

HT29-18N2 cells were differentiated to goblet cells as described previously (24) Briefly, cells were seeded in complete growth medium (DMEM complemented with 10% FCS and 1% P/S), and the following day (Day zero: D-0), the cells were placed in PFHMII protein free medium (GIBCO, ThermoFisher Scientific, Waltham, MA, USA). After 3 days (D-3), medium was replaced with fresh PFHMII medium and cells grown for 3 additional days. At day 6 (D-6) cells were trypsinized and seeded for the respective experiments in complete growth medium followed by incubation in PFHMII medium at day 7 (D-7). All experimental procedures were performed at day 9 (D-9).

### Mucin secretion assay for HT29-18N2 cells

HT29-18N2 cells were differentiated for 6 days and then split into 6-well plates. After one day (D-7), medium was exchanged with fresh PFHMII medium and cells grown for 2 more days. On D-9, cells were washed with isotonic solution containing: 140 mM NaCl, 2.5 mM KCl, 1.2 mM CaCl_2_, 0.5 mM MgCl_2_, 5 mM glucose, and 10 mM HEPES (305 mosmol/l, pH 7.4 adjusted with Tris); and then treated with vehicle (baseline secretion) or 100 μM ATP (stimulated secretion) for 30 min at 37°C. After 30 min at 37°C, extracellular medium was collected and centrifuged for 5 min at 800xg at 4°C. Cells were washed twice in PBS and lysed in 2% Triton X-100/PBS for 1 hr. at 4°C and centrifuged at 14000xg for 15 min. In order to test mucin resistance to shear stress, cells were repetitively washed with isotonic solution after secretion assay and extracellular medium collected.

### Dot blot and WB analysis

Extracellular medium and cell lysates were spotted on nitrocellulose membranes (0.45 μm) using Bio-Dot Microfiltration Apparatus (Bio-Rad, California, USA) (manufacturer’s protocol) and membranes were incubated in blocking solution (5% BSA/0.1% Tween/PBS) for 1 hr. at room temperature. The blocking solution was removed and the membranes were incubated with an anti-MUC5AC antibody diluted 1:2000 or the anti-MUC2 antibody diluted 1:4000 in blocking solution, overnight at 4°C. Membranes were then washed in 0.1% Tween/PBS and incubated with a donkey anti-mouse or anti-rabbit HRP coupled antibody (Life Technologies) for 1 hr. at room temperature. For the detection of ß-actin and FUT8, cell lysates were separated on SDS-PAGE, transferred to nitrocellulose membranes and processed as described for the dot blot analysis using the anti-ß-actin (RRID:AB_476692), anti-FUT8 (RRID:AB_10608850) at a dilution of 1:5000 and 1:1000 in 5% BSA/0.1% Tween/PBS, respectively. Membranes were washed and imaged with LI-COR Odyssey scanner (resolution = 84 μm) (LI-COR, Nebraska, USA). Quantification was performed with ImageJ (FIJI, version 2.0.0-rc-43/1.51 g) (39). The number of experiments was greater than three for each condition, and each experiment was done in triplicates.

### MUC2 and MUC5AC colocalization imaging

Differentiated HT29-18N2 (Control, FUT8 KD or FUT8 OV) cells were grown on 35 mm glass bottom dishes. Next, to visualize MUC5AC and MUC2 colocalization, differentiated HT29-18N2 cells were washed two times, at room temperature, with PBS for 5 min. The cells were then permeabilized by incubation in a buffer (IB) containing 20 mM HEPES pH 7.4, 110 mM KOAc (Potassium acetate), 2 mM MgOAc (Magnesium acetate) and 0.5 mM EGTA (adapted from (40)) with 0.001% digitonin for 5 min on ice, followed by washing for 7 min on ice with the same buffer without detergent. Cells were fixed in 4% paraformaldehyde for 15 min, further permeabilized for 5 min with 0.001% digitonin in IB and blocked with 4% BSA/PBS for 15 min. The anti-MUC5AC antibody was then added to the cells at 1:5000 in 4% BSA/PBS overnight at 4°C; anti-MUC2 antibody was added to the cells at 1:1000 in 4% BSA/PBS overnight at 4°C. After 24 hr., cells were washed with PBS and incubated for 1 hr. at room temperature with a donkey anti-rabbit Alexa Fluor 555 (for MUC2), anti-mouse Alexa Fluor 647 (for MUC5AC) (Life Technologies), diluted at 1:1000 in 4% BSA/PBS, and DAPI (1:20000). Images were acquired using an inverted Leica SP8 confocal microscope with a 63x Plan Apo NA 1.4 objective and analysed using ImageJ ((FIJI, version 2.0.0-rc-43/1.51 g) (39). For detection of the respective fluorescence emission, the following laser lines were applied: DAPI, 405 nm; and Alexa Fluor 555, 561 nm; Alexa Fluor 647, 647 nm. Two-channel colocalization analysis was performed using ImageJ, and the Manders’ correlation coefficient was calculated using the plugin JaCop (41).

### *Fut8*^+/+^, *Fut8*^+/−^, *Fut8*^−/−^ mice

*Fut8*^+/−^ and *Fut8*^−/−^ mice were generated on ICR plus JF1 strain (19), and *Fut8*^+/+^ (Wild type-WT-ICR plus JF1) mice were used as a control for this study (both sets of animals were obtained from Dr. Taniguchi’s and Dr. Gu’s Lab). In order to evaluate the mucus layer under basal situation on healthy mice, 3-month old mice were used (3 females and 1 male). Specifically, four animals per genotype (*Fut8*^+/+^, *Fut8*^+/−^ and *Fut8*^−/−^ mice) were evaluated.

### Histological study

HE stained sections were evaluated under light microscope using a semiquantitative analysis assessing the following parameters: Epithelial hyperplasia (0, normal; 1, minimal; 2, mild; 3, moderate; 4, marked), Inflammatory infiltrate in the lamina propria (0, normal; 1, minimal; 2, mild; 3, moderate; 4, marked), Oedema of the lamina propria (0, normal; 1, minimal; 2, mild; 3, moderate; 4, marked). The total histological score was obtained summing the three parameters (minimum score, 0; maximum score, 12). As an attempt to assess the epithelial regeneration, a counting of the mitotic figures in 20 well-orientated and full-length intestinal crypts was performed. This process was repeated three times and an average was calculated. Other features such as erosion, ulceration, irregular crypts and crypt loss were assessed, but none of the samples showed any of these changes. The histological study was performed in a blinded fashion.

### HE, PAS, PAS-AB, PhoSL, LCA staining

Colon mice sections were stained with haematoxylin and eosin (H/E), periodic acid– Schiff (PAS), Alcian Blue plus periodic acid–Schiff (PAS-AB), PhoSL or LCA for histological analysis. PAS was used to stain neutral, acid-simple non-sulphated and acid-complex sulphated mucins (mucins are stained in purple/magenta). For the PAS-AB staining, it first stains the acidic mucins with Alcian blue; those remaining acidic mucins that are also PAS positive will be chemically blocked and will not react further. Those neutral mucins that are solely PAS positive will subsequently be demonstrated in a contrasting manner. Where mixtures occur, the resultant colour will depend upon the dominant moiety. For PAS-AB staining, acidic mucins are stained in blue, neutral mucins in magenta and mixtures in blue/purple. LCA was used to stain fucosylated proteins, while PhoSL was used to specifically stain FUT8-dependent fucosylation.

Full images of PAS and PAS-AB stained sections were acquired by a NanoZoomer-2.0 HT C9600 scanner (Hamamatsu) at 20X magnification, in which one pixel corresponds to 0.46 μm.

### Measurement of the mucus layer thickness

The thickness of mucus layer was measured in medium and distal colonic tissue sections using the ruler tool of the NDP view + 2.50.19 software (Hamamatsu) in both PAS and PAS-AB stained sections. 20 different measures were performed in two different tissue sections per stain in both medium and distal colon. Mucus layer thickness has been evaluated in all the samples that presented faecal pellets.

### Statistical analysis

All data are means ± SEM. In all cases a D’Agostino– Pearson omnibus normality test was performed before any hypothesis contrast test. Statistical analysis and graphics were performed using GraphPad Prism 6 (RRID:SCR_002798) software. For data that followed normal distributions, we applied either Student’s t test or one-way analysis of variance (ANOVA) followed by Tukey’s post hoc test. For data that did not fit a normal distribution, we used Mann–Whitney’s unpaired t test and nonparametric ANOVA (Kruskal–Wallis) followed by Dunn’s post hoc test. Criteria for a significant statistical difference were: *p<0.05; **p<0.01.

## Acknowledgments

We thank all members of the Malhotra Lab for valuable discussions. We also wish to thank Ms. Noriko Kanto and Ms. Miyuki Kusakawa at the Department of Glyco-Oncology and Medical Biochemistry at Osaka International Cancer Institute for their technical supports. Cell sorting experiments were carried out by the joint CRG/UPF FACS Unit at Parc de Recerca Biomèdica de Barcelona (PRBB). Fluorescence microscopy was performed at the Advanced Light Microscopy Unit at the CRG, Barcelona. V.Malhotra is an Institució Catalana de Recerca i Estudis Avançats professor at the Centre for Genomic Regulation. Histological analysis was performed by Dr. Aguilera and Dr. Prats at the Histology Facility of the Institute for Research in Biomedicine –IRB Barcelona. This work was funded by grants from the Spanish Ministry of Economy and Competitiveness (BFU2013-44188-P to VM), EMBO short-term fellowships (EMBO REF8926 to GCR) and FEDER Funds. We acknowledge support of the Spanish Ministry of Economy and Competitiveness, through the Programmes “Centro de Excelencia Severo Ochoa 2013-2017” (SEV-2012-0208 & SEV-2013-0347) and Maria de Maeztu Units of Excellence in R&D (MDM-2015-0502). This work reflects only the authors’ views, and the EU Community is not liable for any use that may be made of the information contained therein.

## Contributions

GCR and VM conceived and designed the research studies. GCR, CB, YO, TF, YH, AC, NB, GAT and JG conducted the experiments. GCR, YO, GAT, IG, NT and VM analysed and interpreted the data. GCR and VM drafted the manuscript. All authors read, reviewed and approved the final manuscript.

---

**Supplementary Figure 1.** <u>Generation of FUT8 cell lines and their effect on mucin synthesis.</u> **(A)***FUT8* RNA levels normalized to values of HPRT1 from control (Ctrl, black), FUT8 KD (KD, red) and FUT8 OV (OV, green) cells. mRNA levels of each gene are represented as relative value compared to control cells. Results are average values ± SEM (n ≥ 3). **(B)** Cell lysates from control, FUT8 KD and FUT8 OV HT29-18N2 differentiated cells were analysed by western blot with an anti-FUT8 to test expression levels. Actin was used as a loading control. **(C)** MUC1 internal protein levels in differentiated control, FUT8 KD and FUT8 OV cells normalized to the actin levels. Average values ± SEM are plotted as scatter plot with bar graph (n ≥ 3). **(D)** MUC2 internal protein levels in differentiated control, FUT8 KD and FUT8 OV cells normalized to the actin levels. Average values ± SEM are plotted as scatter plot with bar graph (n ≥ 3). **(E)** MUC5AC internal protein levels in differentiated control, FUT8 KD and FUT8 OV cells normalized to the actin levels. Average values ± SEM are plotted as scatter plot with bar graph (n ≥ 3). Abbreviations: Ctrl: Control, m.w.: molecular weight.

**Supplementary Figure 2.** <u>Effect of FUT8 on mucins’ ratio and shear stress resistance.</u> **(A-B)** Relative ratio of MUC2 to MUC5AC secreted levels at baseline (BMS) or after stimulation (SMS) in FUT8 KD (in red) (A) or FUT8 OV (in green) (B) cells and their respective control cells (in black). **(C-D)** Differentiated control (black dots), FUT8 KD (red dots) and FUT8 OV (green dots) cells were incubated for 30 min at 37°C in the absence (Baseline) (C) or presence (Stimulated) (D) of 100 μM ATP. Extracellular medium was collected after successive washes (W0-without washing-, W1 -first wash- and W2 -second wash-) to assess the level of secreted mucins. The y-axis represents MUC5AC secreted levels relative to the total MUC5AC levels for each condition in %. Abbreviations: BMS: Baseline Mucin Secretion, SMS: Stimulated Mucin Secretion, Ctrl: Control. * p<0.05, ** p<0.01.

**Supplementary Figure 3.** <u>Characterisation of *Fut8*^−/−^ colon.</u> **(A)** Histological score of the proximal, middle and distal colon for *Fut8*^+/+^, *Fut8*^+/−^ and *Fut8*^−/−^ mice. Average values ± SEM are plotted as scatter plot, n=4/group. **(B)** Counting of the mitotic figures at the proximal, middle and distal colon for *Fut8*^+/+^, *Fut8*^+/−^ and *Fut8*^−/−^ mice. Average values ± SEM are plotted as scatter plot, n=4/group. **(C)** Representative distal colons of *Fut8*^+/−^ and *Fut8*^−/−^ mice stained with LCA (left panel), PhoSL (central panel) or Hoercht (right panel). **(D)** Ratio of PAS-AB to PAS mucus layer thickness at proximal (P), medial (M) and distal (D) colons of *Fut8*^+/+^, *Fut8*^+/−^ and *Fut8*^−/−^ mice. Abbreviations: ML: Mucus layer. * p<0.05

**Supplementary Table 1.** <u>FUT8 is specifically expressed in colonic goblet cells.</u> **(A)** Data extracted from Tabula Muris (https://tabula-muris.ds.czbiohub.org/).

**Supplementary Table 2.** <u>HE staining’s Histopathological analysis.</u> Abbreviations: Hyp: Epithelial hyperplasia, Inf: inflammatory infiltrate in the lamina propria, Oed: Oedema of the lamina propria, Histo score: Histological score.

