## Supplementary figures and images for "The ulcerative colitis associated gene FUT8 regulates the quantity and quality of secreted mucins"

### Supplemental Figure 1

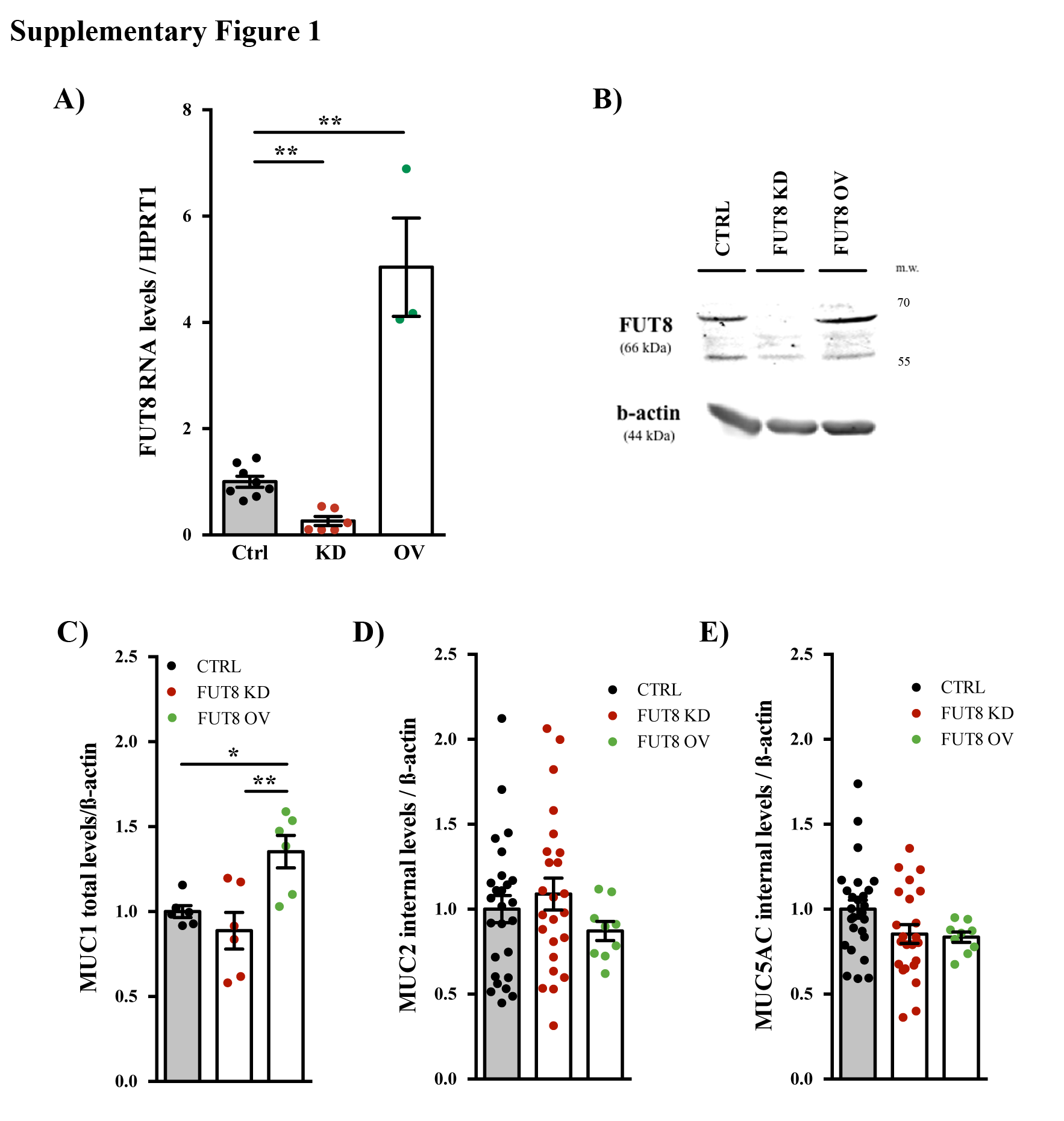

### Supplemental Figure 2

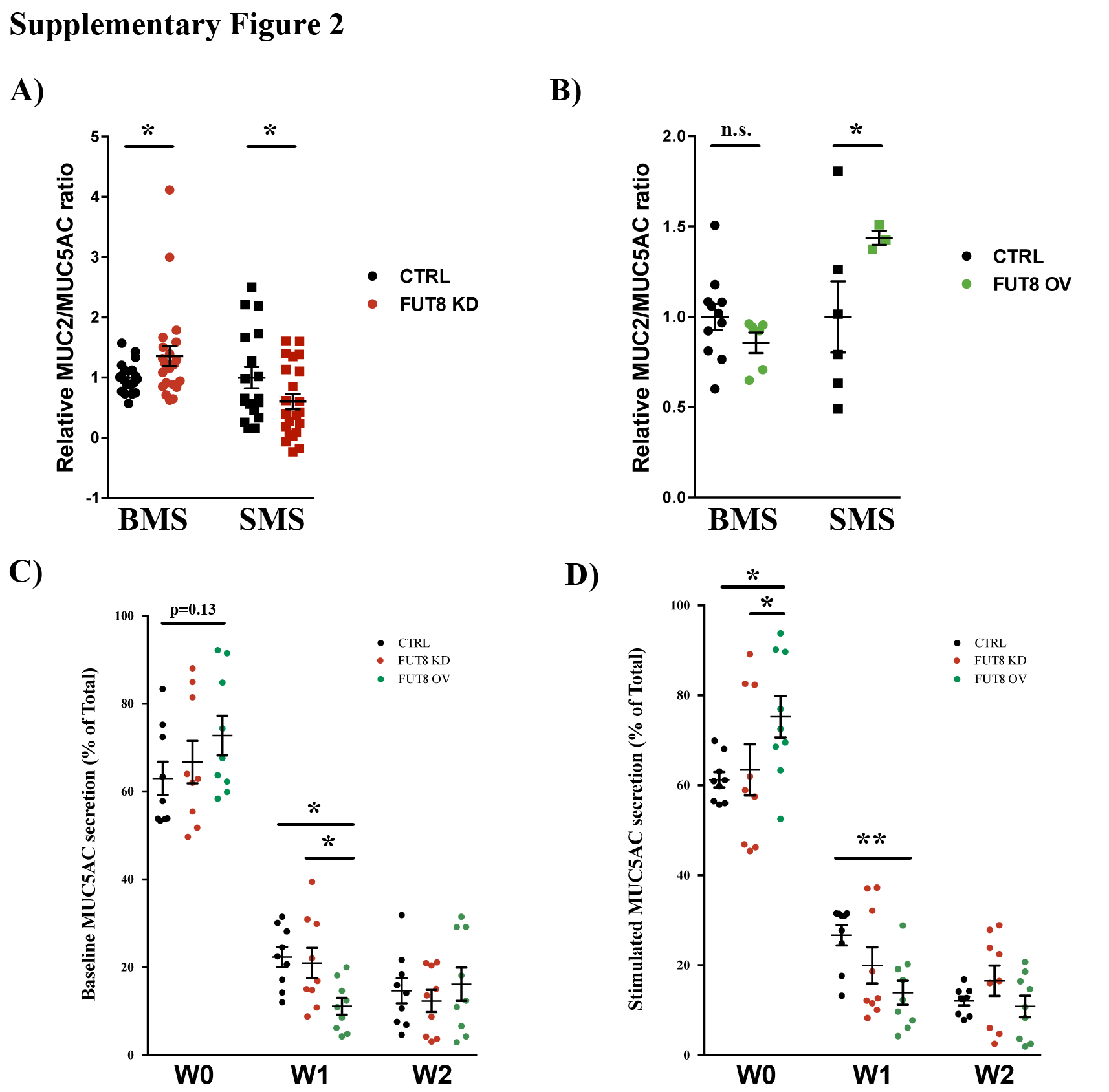

### Supplemental Figure 3

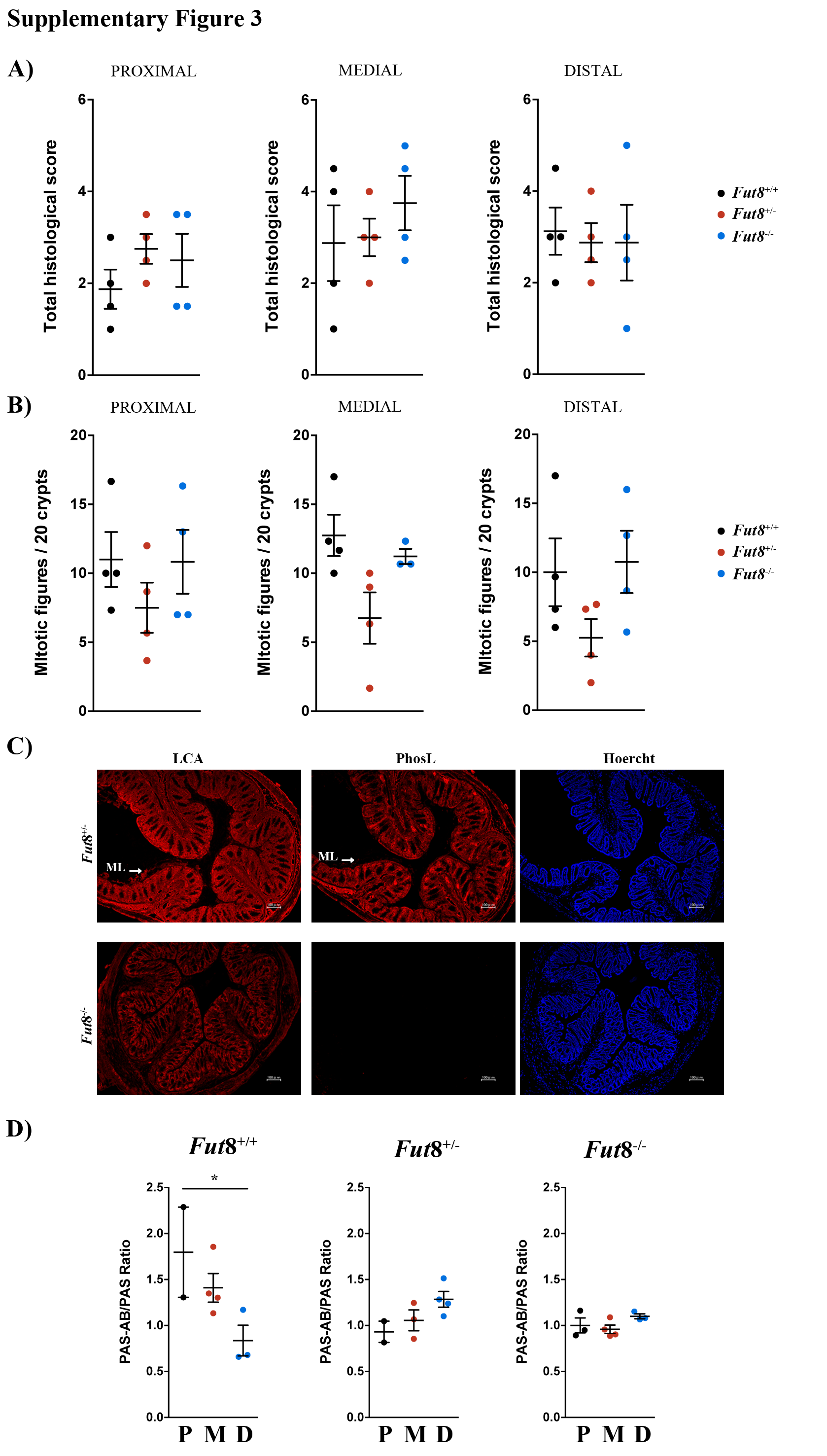

### Supplemental Table 1

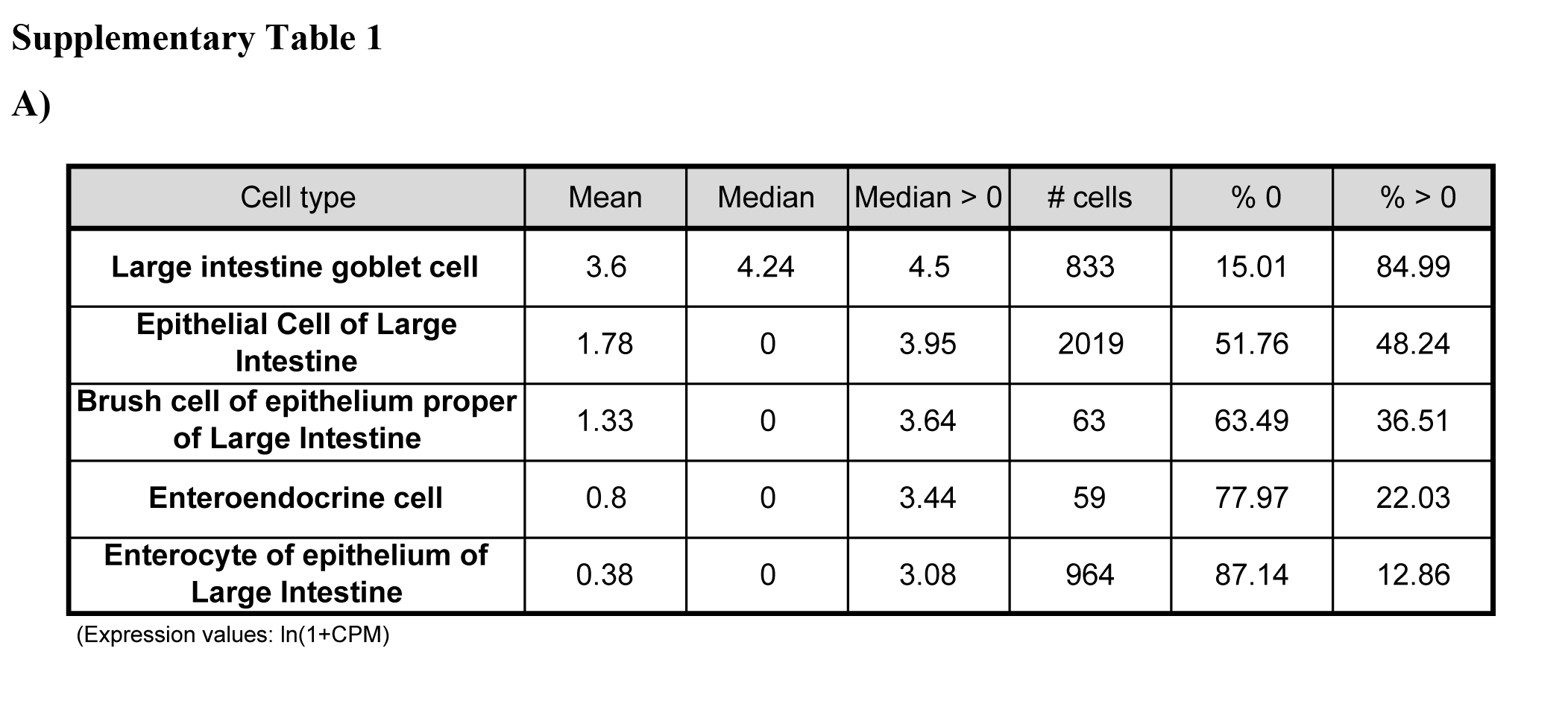

### Supplemental Table 2

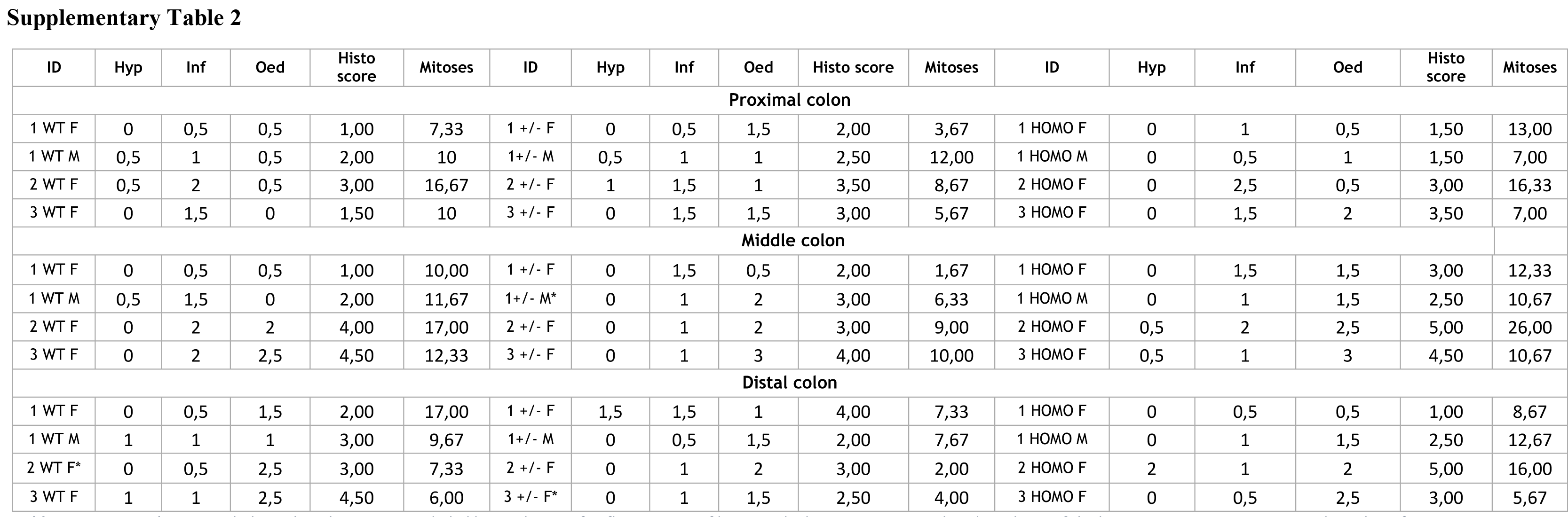
